# Engineering the novel extremophile alga *Chlamydomonas pacifica* for high lipid and high starch production as a path to developing commercially relevant strains

**DOI:** 10.1101/2024.07.18.604193

**Authors:** Abhishek Gupta, João Vitor Dutra Molino, Kathryn MJ Wnuk-Fink, Aaron Bruckbauer, Marissa Tessman, Kalisa Kang, Crisandra J. Diaz, Barbara Saucedo, Ashleyn Malik, Stephen P Mayfield

## Abstract

Microalgae offer a compelling platform for the production of commodity products, due to their superior photosynthetic efficiency, adaptability to non-arable lands and non-potable water, and their capacity to produce a versatile array of bioproducts, including biofuels and biomaterials. However, the scalability of microalgae as a bioresource has been hindered by challenges such as costly biomass production related to vulnerability to pond crashes during large-scale cultivation. This study presents a pipeline for the genetic engineering and pilot-scale production of biodiesel and thermoplastic polyurethane precursors in the extremophile species *Chlamydomonas pacifica*. This extremophile microalga exhibits exceptional resilience to high pH, high salinity, and elevated temperatures. Initially, we evolved this strain to also have a high tolerance to high light intensity through mutagenesis, breeding, and selection. Subsequently, we genetically engineered *C. pacifica* to produce high levels of lipids and starch without compromising growth. We demonstrated the scalability of these engineered strains by cultivating them in pilot-scale raceway ponds and converting the resulting biomass into biodiesel and thermoplastic polyurethanes. This study showcases the complete cycle of transforming a newly discovered species into a commercially relevant commodity production strain. This research underscores the potential of extremophile algae, including *C. pacifica*, as a key species for the burgeoning sustainable bioeconomy, offering a viable path forward in mitigating environmental challenges and supporting global bioproduct demands.

**Graphical Abstract:** 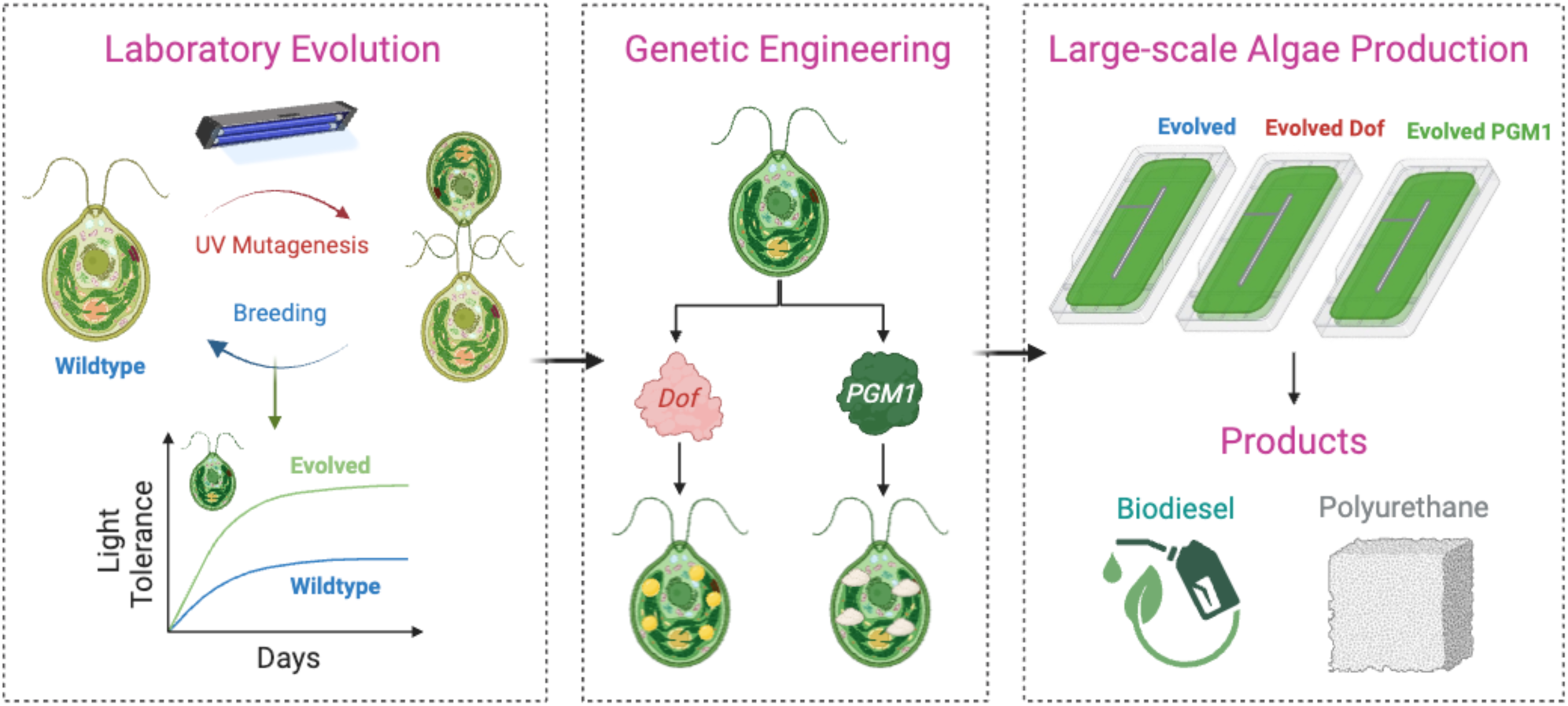

## Introduction

As the 21st century progresses, humanity faces unprecedented challenges in the form of climate change and a rapidly growing global population^1,2^. These twin crises threaten food security, strain conventional energy sources, and exacerbate environmental degradation^3–5^. The quest for sustainable solutions has never been more urgent, compelling the exploration of innovative strategies that can provide the next generation of food and fuel without further harming our planet^6–8^. In this context, developing renewable resources that can reduce carbon emissions and offer a viable alternative to traditional agricultural and energy production methods is critical. The need for such solutions is underscored by the increasing scarcity of arable land, the depletion of freshwater resources, and the necessity to reduce greenhouse gas emissions^9,10^. As we navigate this pivotal moment in human history, the identification and cultivation of sustainable, efficient, and versatile sources of food and fuel emerge not just as a scientific endeavor but as a vital component of our collective response to the existential challenges of our time.

Against this backdrop of environmental and societal challenges, microalgae emerged as a promising solution with the potential to revolutionize our approach to sustainable food, fuel, and material production^11–15^. Microalgae, with their exceptional photosynthetic efficiency, can convert sunlight into biomass more effectively than traditional crops, offering a high-yield, renewable resource that grows rapidly and requires minimal inputs^16^. Their cultivation does not compete for arable land, as they can thrive in non-arable environments, including deserts and saline waters, utilizing non-potable water and even wastewater, thereby reducing the strain on freshwater resources^17^. Furthermore, microalgae are capable of producing a wide range of valuable bioproducts, from biofuels that can reduce our dependence on fossil fuels to high-protein biomass for food and feed, along with bioplastics, pharmaceuticals, and cosmetics^18–22^. This versatility positions microalgae as a potential keystone of a circular bioeconomy, capable of addressing both energy and food security while contributing to carbon mitigation efforts^23^. Harnessing the full capabilities of microalgae as a sustainable resource remains fraught with obstacles. Scaling up production to meet the burgeoning global needs introduces a set of complex technical and financial challenges^24,25^. Chief among these is the necessity to sustain stable cultivation environments on a large scale, a task complicated by the risk of contamination and the variability of environmental factors, which can precipitate culture crashes and disrupt biomass yield^26,27^.

A promising strategy to navigate the hurdles associated with large-scale microalgae cultivation involves focusing on extremophile strains of microalgae^28–31^. These organisms naturally thrive in extreme conditions, such as high salinity, extreme temperatures, acidic or alkaline environments, where conventional microalgae strains would struggle or fail to survive. By leveraging the inherent resilience of these extremophiles, it’s possible to reduce the risk of pond crashes due to environmental stressors, thus ensuring a more stable and reliable biomass production. Furthermore, the genetic engineering of these extremophile strains to enhance their cellular productivity opens new avenues for bioproduct production optimization^32–34^. Through targeted modifications, these strains can be engineered to have increased lipid, carbohydrate, or protein content, thereby maximizing the yield of desired bioproducts. This dual approach of utilizing extremophile microalgae and enhancing their productivity through genetic engineering not only mitigates some of the challenges of stable large-scale cultivation but also paves the way for economically viable and environmentally sustainable production of biofuels, food, and other valuable bioproducts. This innovative strategy harnesses the robustness of extremophiles against environmental challenges while pushing the boundaries of their natural productivity, offering a holistic solution to the sustainability puzzle posed by the increasing demands for food and fuel in the face of climate change.

In this study, we showcase a complete comprehensive pipeline from the genetic engineering of extremophile microalgae in the laboratory to their pilot-scale production for biodiesel and thermoplastic polyurethane synthesis. Our research centers on the novel extremophile species *Chlamydomonas pacifica* (Chlamydomonas Resource Center ID, CC-5697 and CC-5699), which our laboratory discovered in San Diego, California. This microalga is particularly suited for industrial applications due to its exceptional resilience to extreme environmental conditions, such as high pH, high salinity, and elevated temperatures (data unpublished), its ability for mating and high throughput selection, and an ability for recombinant gene expression. Initially, we enhanced the strain’s tolerance to high light intensity through mutagenesis, breeding, and selection. Following this, we employed genetic engineering techniques to develop *C. pacifica* strains that produce high levels of lipids and starch without compromising their growth rates. We successfully demonstrated the scalability of these engineered strains by cultivating them in pilot-scale raceway ponds and converting the resulting biomass into biodiesel and thermoplastic polyurethane (TPU). This study is pioneering in showcasing the complete cycle of transforming a newly discovered species into a commercially relevant strain. The research highlights the potential of *C. pacifica* as a key microalgae species in the sustainable bioeconomy, offering a viable solution to environmental challenges and supporting global demands for renewable resources.

## Results

### Enhancing Light Tolerance through In-vitro Evolution

Given the strain’s natural resilience to high pH, salinity, and temperature, we aimed to identify if we could enhance its tolerance to high light intensity, as well. This improvement would be crucial because high light is known to have an inhibitory effect on microalgae growth^35–38^. To achieve this, we leveraged mutagenesis and breeding approaches, which have been previously shown to improve traits in microalgae^39–42^.

Briefly, we performed UV mutagenesis on the wild-type *C. pacifica* and selected mutants that thrived under high-light conditions (**Figure 1a**, **see Methods**). These selected mutants were mated with another wild-type *C. pacifica,* and progeny were selected under high-light, to identify those strains that showed a stabile resistance to high-light, to ensure the retainment of high-light tolerance trait. Subsequently, the cells underwent another round of UV mutagenesis and selection under high pH tolerance to enhance their resilience to high pH even further. The resultant strain, referred to as the evolved strain, showed remarkable tolerance when exposed to light intensities ranging from 500 µE/m²/s to 3000 µE/m²/s for 24 hours, followed by recovery under low light (80 µE/m²/s) (**Figure 1b**). The evolved strain survived light intensities above 2000 µE/m²/s, whereas the wild-type strain succumbed around 1000 µE/m²/s, indicating a twofold improvement in light tolerance. Moreover, when grown in liquid media, the evolved strain demonstrated approximately a 64% increase in culture density at 750 nm after six days of growth under 400 µE/m²/s and pH 10.25 (**Figure 1d**).

**Figure 1:**
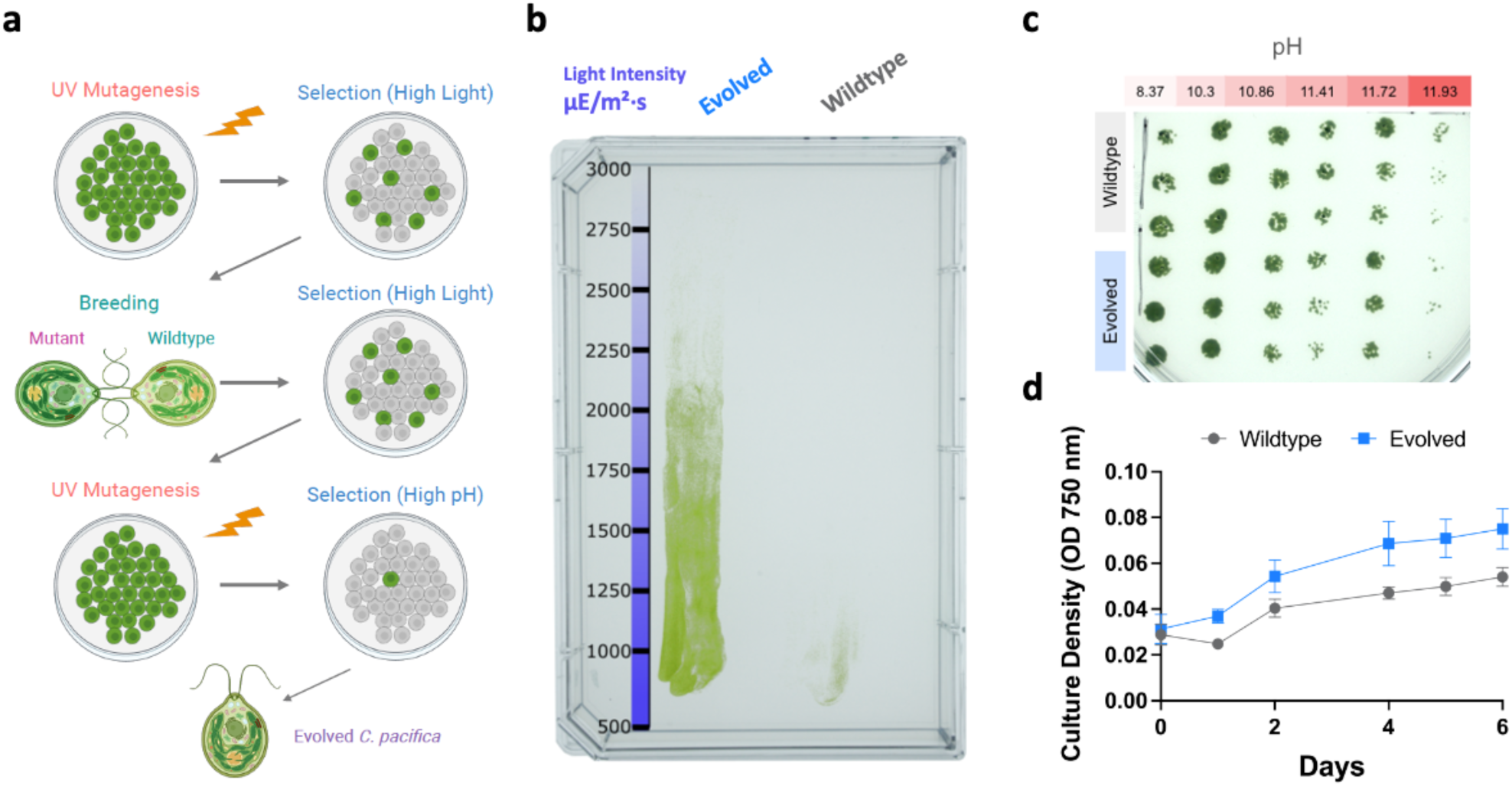
Laboratory evolution of *C. pacifica* for high light tolerance. (**a**) Methodology used for evolving *C. pacifica* to gain high light tolerance, including mutagenesis and breeding. (**b**) Growth comparison of evolved and wildtype *C. pacifica* cells on media plates under a light gradient. (**c**) Tolerance of evolved and wildtype *C. pacifica* cells under different pH conditions. (**d**) Culture density of evolved and wildtype *C. pacifica* cells. Panel (**a**) schematic created with <u>BioRender.com</u>.

Even though we conducted mutagenesis and selection for high pH tolerance in the second round, we could not demonstrate higher pH tolerance between the wild type and evolved strains (**Figure 1c**). This may be due to the cells already achieving their maximum pH tolerance, which remains very high (exceeding pH 11.75), or the loss of the phenotype after subculturing due to the presence of other deleterious mutations in the mutant. A following mating step could be used to stabilize it by reconstituting the deleterious mutations and keeping it beneficial, but this was not pursued at the time^42^. From this point forward, all experiments were conducted using the evolved strain. Overall, these significant improvements highlight the strain’s enhanced adaptability to adverse environmental conditions, validating our approach to genetically refining microalgae for superior resilience to outdoor cultivation. Such advancements not only demonstrate the potential of *C. pacifica* to overcome common cultivation challenges and establish a foundation for advancing large-scale, sustainable microalgae production.

### Genetic Engineering of the Evolved Strain for Enhanced Lipid and Starch Production

#### High Lipid

To enhance the cellular output of the evolved *C. pacifica* strain, we adopted a metabolic engineering strategy centered around transcription factors^33^. Specifically, we overexpressed a soybean-derived *Dof* (DNA binding with one finger) transcription factor, recognized for its effectiveness in boosting lipid production within various plant and microalgae species, including *Arabidopsis*, *Chlamydomonas reinhardtii*, *Chlorella ellipsoidea*, and *Chlorella vulgaris*^43–48^. This family of transcription factors is notable for its heightened responsiveness and greater efficacy in stimulating lipid production, particularly under conditions of nitrogen depletion. We also attempted to identify any endogenous *Dof* transcription factors in *C. pacifica* using iTAK, TAPscan, and PlantTFDB transcription factor identification tools^49–51^. However, none of these tools identified any transcription factors belonging to the *Dof* family in *C. pacifica*. This could be due to the current genome annotation quality limitations for this new species or the evolutionary loss of this transcription factor in *C. pacifica*. Consequently, we proceeded with the exogenous *Dof* transcription factor sourced from the soybean (*Glycine Max*) plant (**Figure 2a, b**).

**Figure 2:**
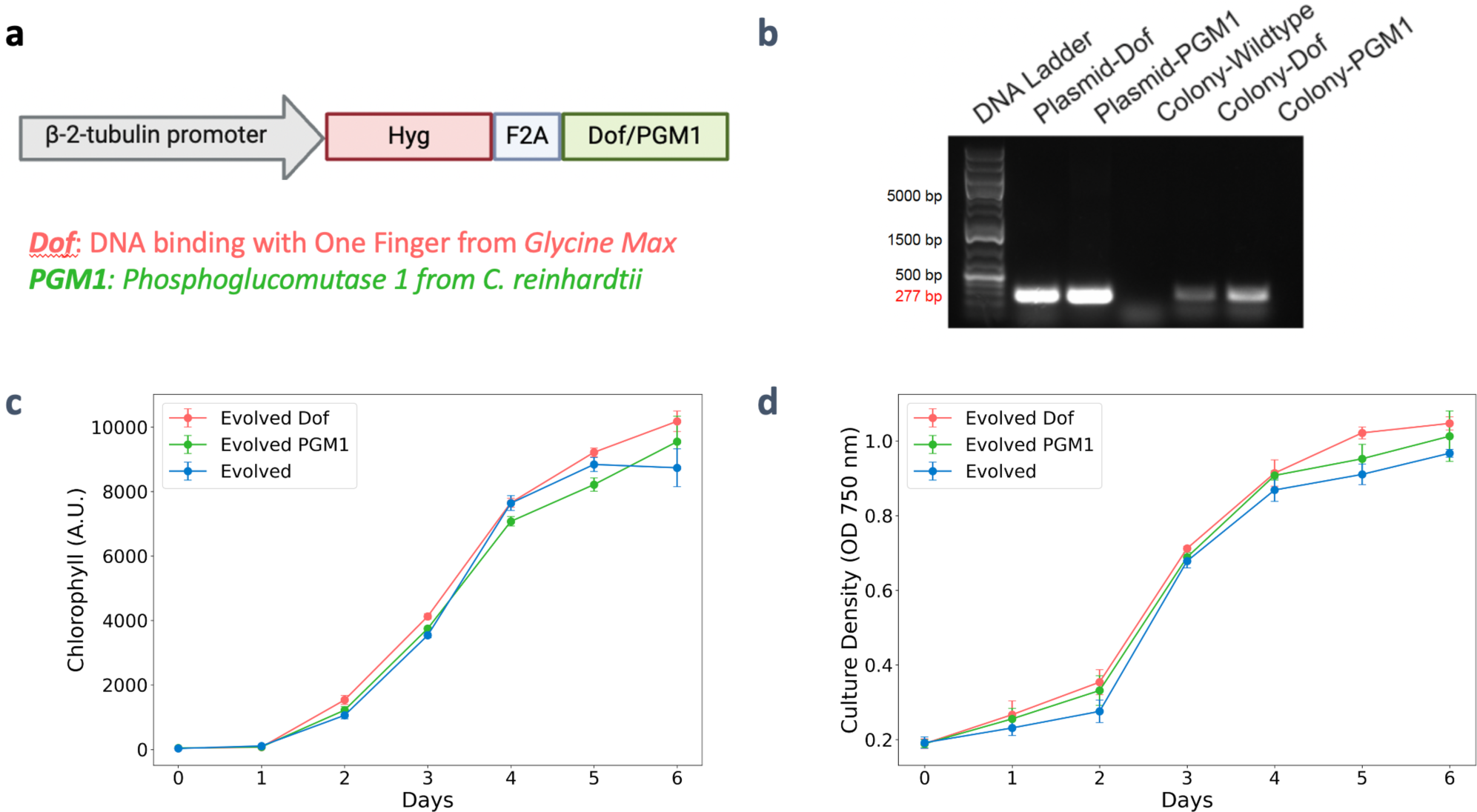
Metabolic engineering of *C. pacifica* for high cellular lipid and starch production. (**a**) Plasmid construct showing β-2-tubulin promoter, target genes, and selectable markers Hygromycin. (**b**) Colony PCR confirmation of transformants. (**c**), (**d**) Growth curves showing chlorophyll and culture density of evolved Dof and evolved PGM1 transformants w.r.t evolved wildtype strain, respectively. F2A represents a self-cleaving sequence. A.U. denotes Arbitrary Unit. Panel (**a**) schematic created with <u>BioRender.com</u>.

Initially, we evaluated the growth performance of the evolved and genetically modified strain (*Evolved Dof*) against the wild-type evolved strain (Evolved) to ensure that overexpression of *Dof* did not negatively impact their growth. Observations indicated no significant difference in culture density at 750 nm and chlorophyll absorption between the transgenic and wild-type strains (**Figure 2c, d**). Additionally, the trait for high-light intensity tolerance remained unchanged by genetic engineering. (**Supplementary Figure 2**). Next, we assessed lipid accumulation in both the evolved and evolved *Dof* strains using Nile Red lipid-binding dye and flow cytometry in the phycoerythrin (PE) channel under both normal and nitrogen-deprived minimal media conditions (**Figure 3a**). To compare the two samples, we measured the population percentage above a threshold of 10^5^ RFU (Relative Fluorescence Units). About 1.2% (± 0.18) of the evolved population in normal media exceeded this threshold, while the Evolved *Dof* population was approximately 29.8% (± 6.62). Under nitrogen-deprived conditions, the evolved population increased to 24.1% (± 4.54), and the Evolved *Dof* population rose to 47.1% (± 5.05). These results indicate that the Evolved *Dof C. pacifica* strain induced more lipid accumulation under normal media conditions compared to the evolved strain, with an even more pronounced effect under nitrogen stress. Confocal microscopy further supported our flow cytometry data, revealing higher lipid-producing cells in the Evolved *Dof* strain compared to the evolved strain under both media conditions (**Figure 3b**). Collectively, these findings demonstrate that current metabolic engineering strategies using transcription factors can effectively enhance cellular lipid productivity in the evolved *C. pacifica* strain.

**Figure 3:**
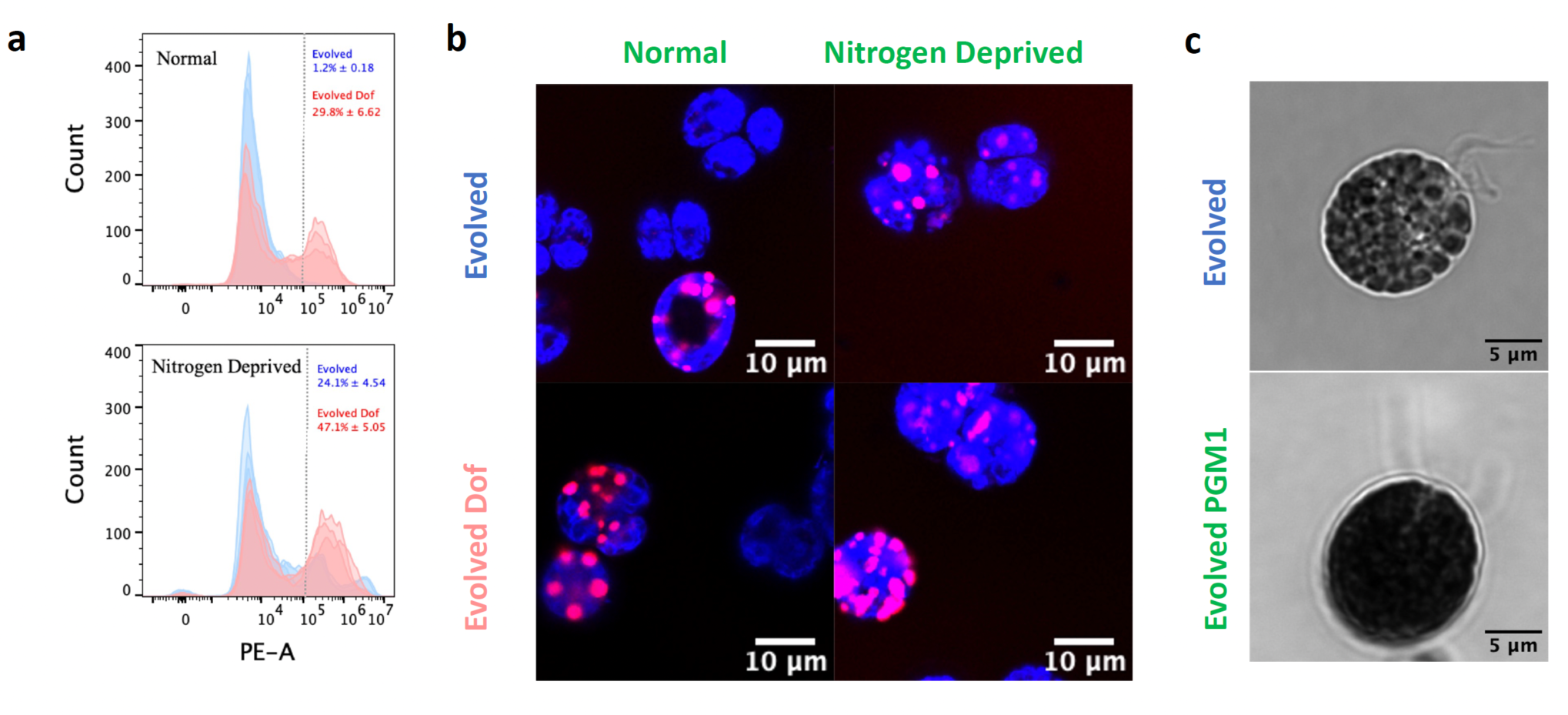
Characterization of evolved *Dof* and evolved PGM1 strains. (**a**) Flow cytometry data of Nile red-stained cells, quantifying lipid content in high lipid-producing *C. Pacifica* cells. (**b**) Confocal microscopy images comparing evolved and evolved *Dof* strains, highlighting lipid accumulation. (**c**) DIC microscopy images of Evolved PGM1 strain compared to Evolved wildtype stained with Lugol solution under sulfur starvation. The gray dotted line in panel (**a**) represents a threshold 10^5^, and PE-A corresponds to the phycoerythrin-area channel. Panel (**a**) shows three curves corresponding to three different biological replicates.

#### High Starch

Phosphoglucomutase (PGM) is an enzyme that catalyzes the interconversion of glucose 1-phosphate (G1P) and glucose 6-phosphate (G6P)^52^. In higher plants such as *Arabidopsis*, *Nicotiana sylvestris*, and *Pisum sativum*, a deficiency in PGM activity results in a starchless phenotype^53–56^. In microalgae, mutants with high starch accumulation have been found to exhibit an increase in phosphoglucomutase 1 (PGM1) expression^57^. These studies indicate that PGM is central in regulating starch accumulation in higher plants and microalgae. Given the pivotal role of PGM in starch accumulation, we overexpressed PGM1 in the evolved *C. pacifica* strain to increase starch accumulation (**Figure 2a, b**). Similar to Evolved and Evolved Dof, Evolved PGM1 retained the trait for high-light intensity tolerance (**Supplementary Figure 2**). We used the Lugol iodine staining method to observe intracellular starch^58^. This method exploits the helical structure of starch, which traps iodine molecules, resulting in a purple-black coloration.

Under Differential Interference Contrast (DIC) microscopy, we observed a higher intracellular black coloration in the evolved PGM1 cells compared to the evolved strain under sulfur starvation (**Figure 3c**). Additionally, in the plate reading assay, we noted a 30% higher normalized absorbance at a wavelength of 660 nm (**Supplementary Figure 3**). Altogether, these results indicate that the PGM1 overexpressed evolved strain accumulates more starch than the evolved strain.

#### Pilot-scale cultivation of high-lipid and high-starch evolved strains

Scaling lab strains to larger-scale production in ponds often results in culture failures due to operational issues, human error, biological contamination, or environmental factors. These challenges pose significant barriers to transitioning lab strains to industrial-scale applications^59–62^.

To assess the scalability and commercial viability of our genetically engineered high-lipid and high-starch evolved *C. pacifica* strains, we initiated pilot-scale cultivation trials of the evolved, evolved *Dof* and evolved PGM1 transgenic strains in 80-liter open raceway ponds within a greenhouse environment (**Figure 4a**). We began by culturing strains in 250 ml flasks and then scaling these up to 20-liter carboys, still in lab settings (see **Methods**). The biomass from the carboys was then used to inoculate the 80-liter ponds, which were operated photoautotrophically. We conducted three consecutive rounds of biomass production in these ponds in San Diego, California, during November and December inside a closed greenhouse and monitored growth conditions such as light intensity and air temperature. The ponds shown in **Figure 4a** are from the end of round 1 growth. **Figures 4b** and **4c** display the average daily light (measured in PPFD, photosynthetic photon flux density) and average daily temperature (measured in degrees Celsius), respectively. Remarkably, given the greenhouse effect, there were days when the temperature inside the greenhouse reached up to 50°C, demonstrating the resilience of these strains in a more natural and unconstrained setting.

**Figure 4:**
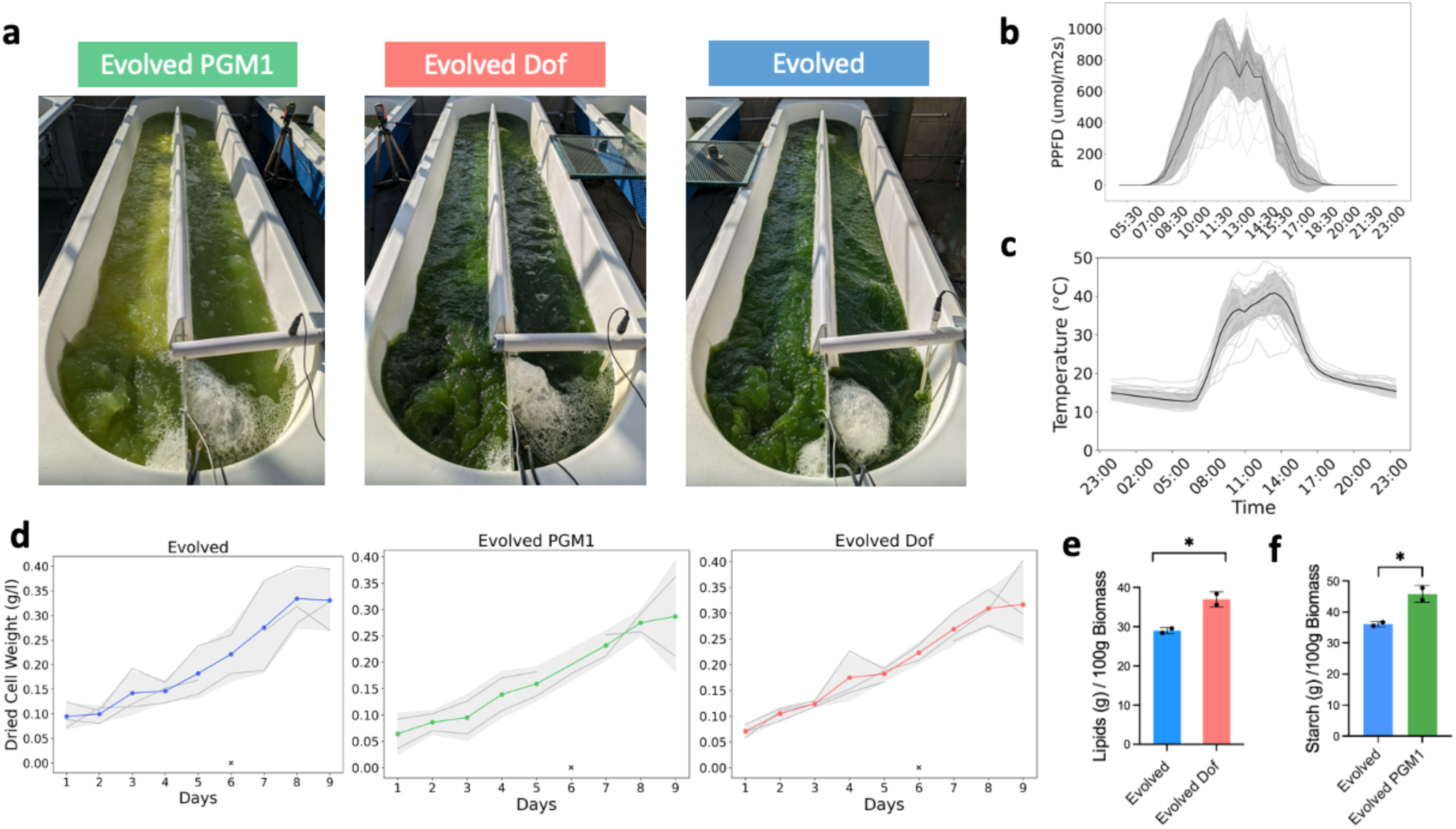
Pilot-scale production of high lipid and high starch strains. (**a**) Visuals of raceway ponds showcasing cultivation of evolved and engineered *C. Pacifica* strains. (**b**), (**c**) Light sensor and temperature sensor data recorded during the pond runs, respectively. (**d**) Dry cell weight (g/l) measurements throughout three consecutive pond runs. (**e**), (**f**) Lipid (g) and Starch (g) measurements relative to dried biomass (g), respectively. In panel (**b**), PPFD represents Photosynthetic Photon Flux Density, and in panels (**b**) and (**c**), a break in the line represents loss of sensor data. In panel (**d**), “x” corresponds to the day of partially missing data. In panel (**e**), * denotes p < 0.05. The p-values were calculated using an unpaired t-test.

We also measured the dried cell weight (g/l) of the three strains to assess biomass productivity, which reached up to 0.4 g/l in 8-9 days (**Figure 4d**). Interestingly, we observed stagnation or no growth in the PGM1 strain in the first round. This might be due to human error or potential contamination from the inoculum prepared in minimal media supplemented with acetate as the carbon source (**Figure 4a**). Despite this initial challenge, the PGM1 strain exhibited a remarkable recovery in subsequent rounds, achieving growth and productivity comparable to the wildtype and the *Dof* strain, underscoring its resilience and adaptability.

We also performed gravimetric analysis on lipids and starch extracted from biomass grown in ponds to determine if the high-lipid and high-starch phenotype is consistent at scale. In the evolved Dof strain, lipid content was approximately 28% higher (p = 0.032, unpaired t-test) compared to the evolved strain, with lipid content values of 28.99 g ± 0.78 / 100 g dried biomass and 37.00 g ± 1.93 / 100g dried biomass, respectively (**Figure 4e**). Moreover, we observed a 27% increase in starch content (p = 0.039, unpaired t-test; evolved: 36.08 g ± 0.8 / 100g dried biomass, evolved PGM1: 45.83 g ± 2.68 / 100g dried biomass) in the evolved PGM1 strain compared to the evolved strain (**Figure 4f**).

Altogether, these findings collectively affirm the potential of the engineered *C. pacifica* strains to thrive in large-scale operations. Their demonstrated resilience to extreme environmental conditions, coupled with their recovery and sustained productivity, positions these strains as promising candidates for commercial-scale cultivation, offering a tangible pathway toward the production of sustainable bioproducts.

### Bioproducts

#### Biodiesel

Oleaginous microalgae have been targeted as a promising feedstock for the production of renewable biodiesel. However, challenges in scalability and product purity associated with lipid extraction and chemical conversions into fuel have limited this application thus far^63^. Traditional approaches for renewable biodiesel production begin with lipid extraction and recovery from biomass to generate a total lipid extract that can be chemically converted to fatty acid methyl esters. Still, lipid extraction methods pose challenges in efficiency and efficacy^64^. Additionally, while microalgae produce high amounts of neutral lipid triglycerides (TAGs) that can be converted into biodiesel, other classes of lipids, including nonpolar sterols and polar glycolipids, are present in total lipid extracts and can act as pervasive contaminants during biodiesel synthesis^64^.

To avoid the difficulties associated with traditional lipid extraction-chemical conversion methods of biodiesel production, we have successfully used a two-step direct-saponification esterification method to afford high-purity biodiesel directly from algal biomass^65^. In this process, dried biomass is treated with basic sodium hydroxide to selectively convert neutral triglycerides and free fatty acids within the algal cell into water-soluble charged sodium soaps (**Figure 5a**). This aqueous solution is extracted with hexane solvent to remove remaining uncharged lipid contaminants (nonpolar and polar lipids), enabling selective isolation of relevant biodiesel precursors. Treatment of the aqueous layer with hydrochloric acid to pH<5 serves to reprotonate fatty acid soaps and form free fatty acids, which can then be easily recovered in an organic solvent. An aliquot of free fatty acids was saved for further characterization, and the remaining sample was converted into fatty acid methyl ester (FAME) biodiesel with methanol and sulfuric acid (**Figure 5a**). All four samples, including evolved and evolved *Dof* under minimal and nitrogen-deprived media, of *C. pacifica* produced yellow liquid biodiesel ranging from 3-12% of the initial cellular dry weight.

**Figure 5:**
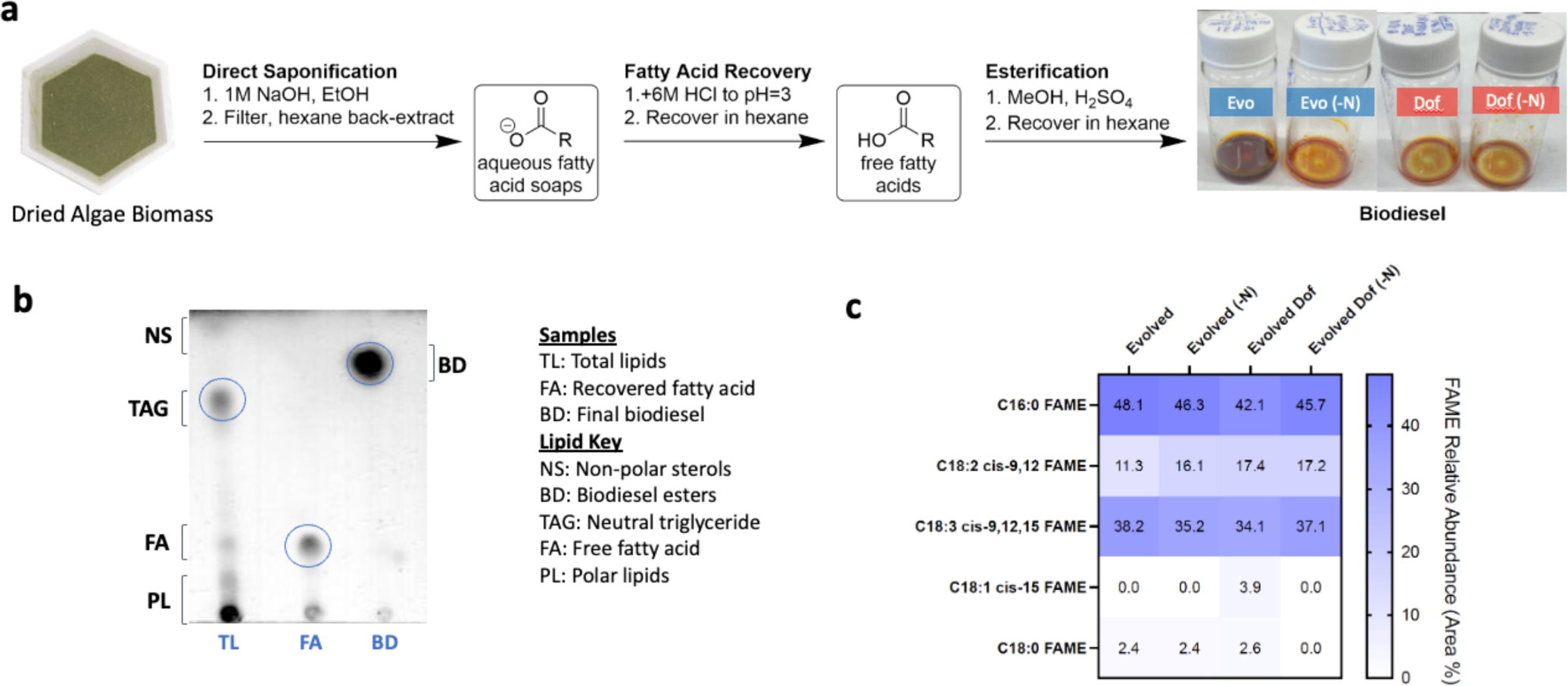
Conversion of algal biomass to biodiesel. (**a**) Scheme of the conversion process from algal biomass to biodiesel. (**b**) Representative TLC analysis of biodiesel production process (Dof sample shown). (**c**) Heatmap showing GC-MS analysis of biodiesel content (n=1). In panels (**a**) and (**c**), the nitrogen-deprivation condition is denoted with (-N); otherwise, the samples were in normal minimal media. In panel (**c**), FAME stands for Fatty Acid Methyl Esters.

The purity of intermediate free fatty acids and final biodiesel was monitored using thin layer chromatography (TLC) with a specialized eluent and staining system that can visualize different lipid classes^66^. An example TLC of the *Dof* intermediate and product is shown in **Figure 5b**. When compared to a standard of total lipids extracted from *C. pacifica*, the intermediate free fatty acids and final biodiesel products displayed high purity, exemplified by the presence of only one major product spot in the TLC. The biodiesel sample showed a significant product with a retention factor (Rf) value of 0.83 compared to the fatty acid spot with an Rf of 0.25 and the TAG spot seen in the total lipids with an Rf of 0.70. This aligned with the expectations that the biodiesel product would be less polar than the initial TAGs and fatty acids and thus have a higher retention factor through TLC.

To confirm the successful synthesis of biodiesel, gas chromatography-mass spectrometry (GC-MS) was used to analyze and identify the components of each sample. Final biodiesel samples were diluted into hexane for injection, and the GC retention times and MS mass-to-charge peaks were used to identify methyl ester biodiesel products. Products from all four algal cultures displayed high purity (>80%) of mixed methyl ester biodiesel. The primary component of these mixed biodiesel samples was hexadecenoic acid methyl ester (C16:0 FAME), ranging from 42-48% relative abundance compared to the FAME mixture (**Figure 5c**). Other saturated (methyl stearate, C18:0 FAME) and unsaturated (C18:2 cis-9,12 FAME, C18:3 cis-9,12,15 FAME) biodiesel components were also observed in all samples. Overall, the ease of this method for producing microalgal biodiesel combined with the high purity seen in end products demonstrates the promising capability of *C. pacifica* as a producer strain of high-value industrial products.

#### Thermoplastic polyurethane

To showcase the potential of the *C. pacifica* strain in practical applications, we successfully formulated thermoplastic polyurethanes (TPUs) using downstream processing of its algal biomass. We chose to design TPUs for two main reasons. First, the inherent production capabilities of the strain to produce starch and fatty acids, both of which can be converted into useful diol, diacid, and diisocyanate monomers – essential building blocks for polyurethanes^67^. Secondly, polyurethanes account for roughly 10% of global plastic production and are ubiquitous in our everyday lives, finding applications in coatings, adhesives, foams, and more^68^.

The synthesis of an algae-based TPU, designated as A2141, was achieved using *C. pacifica* as the main renewable source (**Figure 6a, b**). First, the synthesis of an algal polyester polyol was prepared via a polycondensation reaction, as previously reported^69,70^. The resultant polyol had a molecular weight of 1261 g/mol by hydroxyl titration and a polydispersity index (PDI) of 1.68 by gel permeation chromatography (GPC). The algal polyol was reacted with an algae-derived aliphatic diisocyanate and cured in an oven at 80° for 3 days. Once cured, the TPU had a molecular weight of 91 kDa and a PDI of 2.58 by GPC. A2141 exhibited a shore A hardness of 75 ± 3, tensile strength of 3.22 MPa, and an elongation of 282% at break.

**Figure 6:**
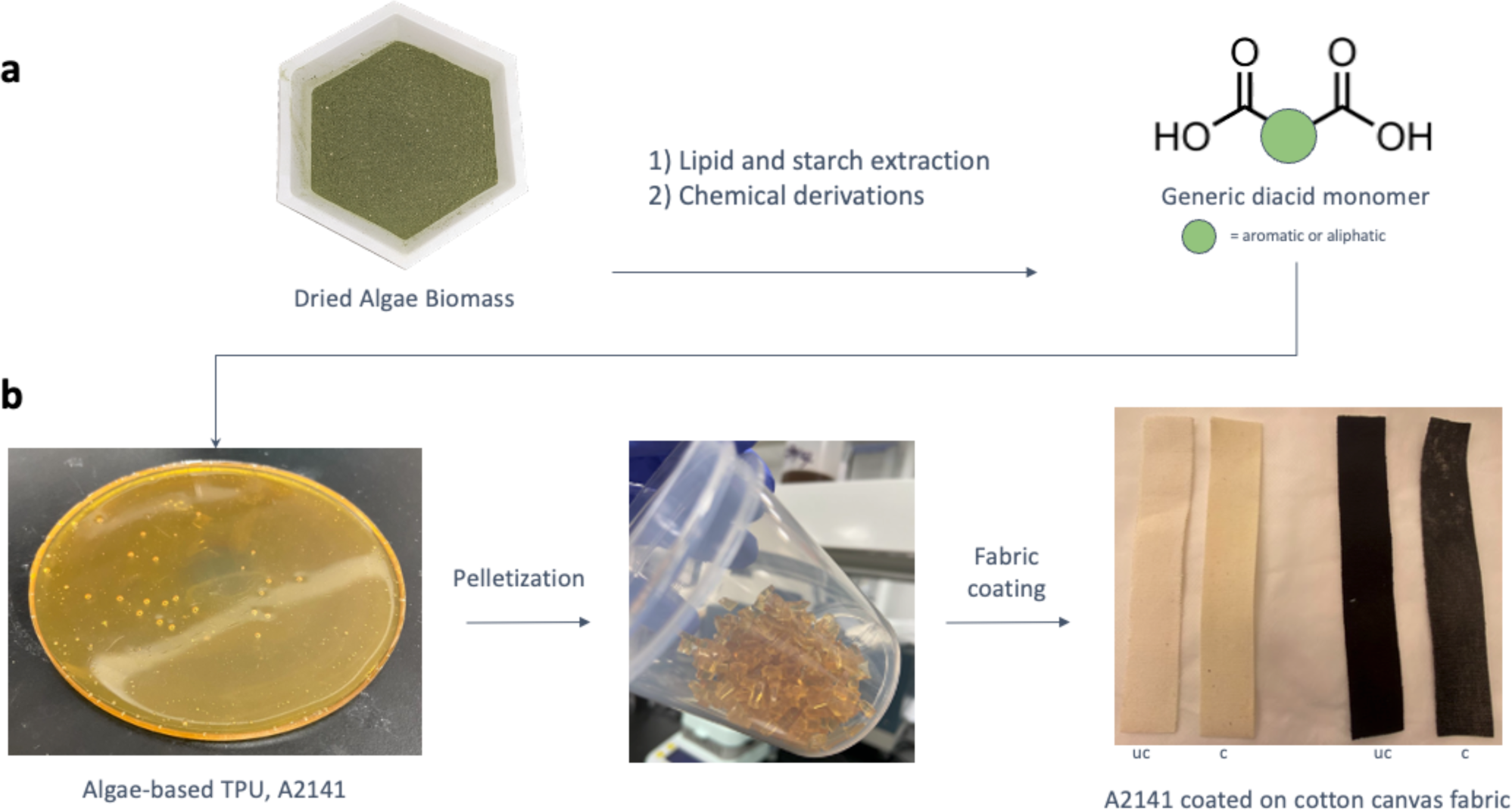
Conversion of algal biomass to TPU-coated fabric. (**a**) Conversion of algal biomass to polyurethane (PU) foam. (**b**) Thermoplastic polyurethane, A2141, coated on cotton canvas. (uc: uncoated, c: coated)

To demonstrate possible applications, A2141 was coated onto cotton canvas fabric with a set thickness of 0.05mm using a solution of 18% A2141 in dimethylformamide (DMF). The coated cotton fabric displayed hydrophobic characteristics with a water contact angle of 97.99° ± 0.5°, which is greater than that of polyethylene terephthalate (PET) commonly found in water bottles, and polypropylene (PP) used in the packaging industry, which has contact angles of 91.3° and 105°, respectively^71,72^. The material stiffness, or flexural rigidity (G), was also analyzed by comparing the flexibility of the uncoated cotton and the algae-TPU-coated cotton. We observed a 12-fold increase in flexural rigidity after the coating treatment. The flexural rigidity was determined to be 329 mg*cm for the uncoated fabric and 4058 mg*cm for the coated fabric, thus reinforcing the fabric and contributing to the hydrophobicity of the material.

These results indicate that a 75% algae-based material could have significant commercial relevance in applications such as the fashion industry, particularly for waterproofing fabrics used in raincoats and duffle bags. Additionally, they highlight the importance of engineering robust algae strains for plastics production.

## Discussion

The potential of microalgae as biofactories has already been demonstrated by their ability to produce a diverse range of bioproducts, including biofuels, nutraceuticals, and bio-based chemicals^11–15,33^. However, large-scale cultivation of microalgae continues to present significant challenges, primarily due to uncontrolled growth conditions, the risk of contamination, and pond crashes^26^. Our lab recently discovered a novel extremophile, *C. pacifica*, demonstrating remarkable growth resilience under high pH, high temperature, and high salinity conditions. Through mutagenesis and breeding, we have further evolved this strain to tolerate high light intensity, making it a robust candidate for open pond cultivation.

*C. pacifica*, belonging to the same genus as the well-studied *C. reinhardtii*, allows for the opportunity to leverage existing research on *C. reinhardtii* to accelerate the development of *C. pacifica* as an industrial strain. Methods to enhance bioproduct productivity in *C. reinhardtii* can be directly applied *to C. pacifica*. In this study, we improved lipid and starch content in evolved *C. pacifica* by overexpressing the soybean *Dof* transcription factor and the PGM1 gene from *C. reinhardtii* and demonstrated significant product enhancements (**Figure 3, 4**). Similar genetic engineering strategies, such as overexpressing chloroplast-type glyceraldehyde-3-phosphate dehydrogenase (cGAPDH) to enhance overall biomass productivity or utilizing hybrid breeding approaches with selection-enriched genomic loci (SEGL) to boost photosynthetic productivity under diverse conditions could be used to further streamline the development of *C. pacifica* into a robust, high-yield strain^73,74^.

Here, we successfully demonstrated that the evolved *C. pacifica* can be grown in pilot-scale raceway ponds, leveraging its pH tolerance to avoid contamination and crashes (**Figure 4**). While the evolved wildtype and evolved *Dof* strains grew well without any crashes in three consecutive rounds, the evolved PGM1 strain experienced a crash in the first round but recovered in the subsequent rounds. We suspect the initial crash was due to contamination of the inoculum, prepared under heterotrophic conditions in a medium with acetate as the carbon source at a nearly neutral pH. To mitigate any future contamination risks, preparing the inoculum at a higher pH and using CO_2_ captured through direct air capture technologies could be valuable^75^. This approach may reduce contamination risk and potentially make the process a net-positive source for the bioeconomy. However, further life cycle assessment is required to validate this cultivation method^76^. Moreover, given the growing interest in sustainable aviation fuel (SAF) within the airline industry, it would be highly beneficial to test the conversion of lipids extracted from the engineered strain of *C. pacifica* into SAF^77,78^. Additionally, performing a techno-economic analysis would provide a comprehensive understanding of *C. pacifica’s* potential as a viable biofactory for producing SAF.

Most studies have focused on extracting a single valuable feedstock, such as lipids, carbohydrates, or proteins, from algal biomass. To enhance the economic viability of algae, it would be beneficial to optimize downstream extraction processes to efficiently extract all proteins, carbohydrates, and lipids simultaneously^79,80^. These can then be converted into valuable products such as therapeutic proteins, animal feed, biomaterials, and biofuels. This comprehensive approach would reduce the burden and high cost associated with a single commodity and help integrate algae into the mainstream of biotechnology. *C. pacifica* is an excellent candidate for such tests, given its compatibility with genetic engineering and large-scale cultivation.

## Conclusion

This study demonstrates the potential of developing extremophile microalgae as a commercially viable production platform. Here, we have demonstrated the successful transformation of the novel extremophile species, *C. pacifica*, into a robust, high-yield strain. Additionally, pilot-scale cultivation has shown the feasibility of producing bio-based fuel and material, highlighting the potential for large-scale applications. By enhancing its resilience, employing genetic engineering, and achieving large-scale growth, we have demonstrated *C. pacifica’s* potential to produce valuable bioproducts such as biofuels and polyurethane. The versatility of *C. pacifica* as a biofactory highlights the promise of extremophile algae in a future sustainable bioeconomy, offering potential solutions to address climate change and global resource demands. Future research should focus on optimizing downstream extraction processes and conducting comprehensive techno-economic analyses to fully realize the potential of *C. pacifica* for industrial applications.

## Method

### Media and algae strain

For lab flask cultures and 20-liter carboys, High-Salt Media with acetate (HA) was used, with or without high pH. If the media had a high pH, NH₄Cl was replaced with urea, and the pH was raised to 10.5 to avoid the release of NH₃. For nitrogen-deprived media, the nitrogen source (NH₄Cl or urea) was removed. Ammonium nitrate was used as a nitrogen source for pond pilot runs. For pond main runs, urea was used as the nitrogen source, and the pH was increased and maintained daily at 10.5 using NaOH. The medium contained 40-100 mM NaCl and no additional carbon source, thus operating under fully phototrophically conditions.

*C. pacifica* (CC-5699) and *Evolved C. pacifica* (CC-6190 402wt x 403wt) were used throughout all experiments, which can be ordered from Chlamydomonas Resource Center at the University of Minnesota in St. Paul, Minnesota. In the lab, the cells were grown under continuous 24-hour light conditions at approximately 25°C, with a photon flux of 125 µE/m²/s, on shaken tables rotating at 125 RPM.

### Mutagenesis and breeding

To generate a killing curve, cultures in petri dishes (150 mm) were prepared at a cell density of 1-2 × 10^7^ cells in the stationary phase. The cells were grown in HA medium and exposed to UV light using BioRad Gs Gene Linker UV Chamber, with exposure intervals of 5 seconds. Samples were taken between exposures and dotted onto HA agar plates. These plates were incubated for 7 days, and survival was evaluated to determine the exposure duration at which noticeable cell death occurred, but some viable cells remained, identified to be between 35-40 seconds.

For mutant library generation, cells grown to the stationary phase were exposed to 35-40 seconds of UV light under the same conditions as the killing curve experiment. Following UV exposure, cells were allowed to recover for 24 hours in fresh media. During the screening phase, cells were plated on a high-light screening plate. Cells at the edge of tolerance were scrapped, recovered, and grown again until reaching the stationary phase. These cells were then bred with the parental cells to enhance the desirable traits^81^.

### High light tolerance assay

The strains were exposed to high light conditions, ranging from a minimum of 500 µE/m²/s to a maximum of 3000 µE/m²/s, over a continuous 24-hour period, followed by incubation at 80 µE/m²/s for five days. The gradient of light exposure is visually represented by the decreasing density of algal growth from the bottom (500 µE/m²/s) to the top (3000 µE/m²/s) of each plate. The gradient was generated by adding a strong light source (HERB PRO G2, 1000W Mastiff Growl LED Grow Light) with blue and red LEDs 9 cm from the plates. Positioning the plates on the corner of the panel allows for a linear gradient light intensity measured with a photometer (**Supplementary Figure 1**).

### High pH tolerance assay

The cells were grown up to late exponential to stationary phase and were transferred to high pH media for 24 hours. After 24 hours, cells were plated and grown for 6-7 days.

### Vector design and algae transformation

The vector was constructed using genetic elements derived from the assembled genome of *C. pacifica.* The β-Tubulin A 2 gene promoter was utilized to drive the expression of hygromycin (used as a selection marker) and the *Dof* or *PGM1* gene. The complete map can be found at Zenodo (https://doi.org/10.5281/zenodo.12636981). The Dof (pJPCHx1_Dof_1.1) and PGM1 (pJPCHx1_PGM1) vectors have been submitted to the Chlamydomonas Resource Center at the University of Minnesota in St. Paul, Minnesota.

For the transformation process in algae, plasmid DNA was digested using KpnI and XbaI enzymes (New England Biolabs, Ipswich, MA, USA). This was followed by purification with the Wizard SV Gel and PCR Clean-up System (Promega Corporation, Madison, WI, USA) without separating the fragments. DNA concentration was measured using the Qubit dsDNA High Sensitivity Kit (Thermo Fisher Scientific, Waltham, MA, USA). Electroporation was used for transformation, following the method described by Molino et al., 2018^82^. Transformed cells were selected on HA agar plates containing 30 µg/mL hygromycin.

The transformants were verified through Colony-PCR using primers specific for a 277 base pair segment of the hygromycin gene (forward primer: TGATTCCTACGCGAGCCTGC; reverse primer: AACAGCTTGATCACCGGGCC), followed by sequencing.

### Plate reader assay

A volume of 160 µL from each sample was placed in a flat-bottom 96-well plate (Corning Costar, Tewksbury, MA, USA). Fluorescence was measured using a Tecan Infinite M200 Pro plate reader (Tecan Group Ltd., Männedorf, Zurich, Switzerland). Chlorophyll fluorescence was recorded with an excitation wavelength of 440 nm and an emission wavelength of 680 nm, while absorbance was measured at a wavelength of 750 nm.

### Nile red staining and flow cytometry

A 50 µL aliquot of culture was diluted with 110 µL of buffer to reach a final volume of 160 µL. Then, 1.6 µL of 1 mM Nile red in DMSO was added to the solution. The cells were incubated at 40°C for 10 minutes using the Tecan Infinite M200 Pro plate reader (Tecan Group Ltd., Männedorf, Zurich, Switzerland). The stained and incubated samples were analyzed using a Beckman Coulter CytoFLEX Cytometer, utilizing the PE channel (585/42 bandpass filter) to detect Nile Red fluorescence (excitation/emission 551/636 nm) bound to intracellular lipids with a gain setting of 1. 10,000 events were recorded at a 30 µL/min flow rate for each sample.

### Microscopy

For Differential Interference Contrast (DIC) microscopy, cells were stained with 1% Lugol’s solution and imaged using a Nikon Eclipse Ni microscope. Images were taken at 60X magnification plus the ocular lens and the same gain and exposure settings were maintained for all samples to ensure consistency.

For confocal microscopy, sample slides were prepared using Frame-Seal™ Slide Chambers containing agar supplemented with media following the protocol described in dx.doi.org/10.17504/protocols.io.bkn8kvhw. Cells were placed on the agar and sealed with a glass cover slip. Confocal microscopy was performed at the UCSD School of Medicine Microscopy Core using a Leica STED microscope. An excitation wavelength of 405 nm with an emission range of 630-680 nm was used for chlorophyll detection. An excitation wavelength of 470 nm with an emission range of 475-575 nm was utilized for Nile red. The emission and excitation settings were carefully adjusted to minimize channel cross-talk.

### Raceway Ponds

Three 80-liter raceway ponds were utilized for algae cultivation inside a greenhouse. The ponds were filled with HSM prepared using normal tap water. For pilot runs, we used ammonium nitrate as the nitrogen source. However, urea served as the nitrogen source for later main runs, and the pH was elevated to 10.5 using NaOH. The pH was monitored and maintained at 10.5 daily. The media contained 40 mM NaCl and excluded any additional carbon source, enabling fully phototropic growth. The ponds were exposed to natural sunlight, and algal growth was monitored daily by measuring chlorophyll (440/680), absorbance at 750 nm, and dry cell weight (DCW, g/L). For DCW measurement, a 50 ml sample was collected, salts were removed using EDTA, and the sample was dried in an oven to determine the dried biomass weight. The greenhouse was equipped with sensors to monitor culture pH, culture temperature, light intensity, and air temperature.

### Lipid Extraction

Lipid extraction was carried out on the evolved and evolved Dof samples from the pilot run, each having two technical replicates. A 50 ml culture was centrifuged and dried. The dried biomass was then subjected to the Bligh Dyer lipid extraction method^83^. The extracted lipids were weighed, and the data was presented as a percentage of the dried biomass.

### Starch Extraction

Following lipid extraction, the residual biomass was subjected to several cycles of extraction using 80% ethanol and 50 mM NaOH to remove any remaining protein and lipid content. This process was repeated until a white sediment was observed. The sediment was then washed twice with water to remove any residual ethanolic solution and subsequently dried. The initial dry cell weight was approximately 100 mg, and the recovered dried material was weighed to quantify the extracted starch.

### Conversion of algal biomass to biodiesel

#### Biodiesel synthesis

Algal biodiesel was prepared via direct biomass saponification followed by acid-catalyzed Fischer esterification^84^. Wet algal biomass slurry samples were flash frozen in a dry ice/isopropanol bath prior to lyophilization to sublimate excess aqueous media (LabConco FreeZone 2.5, 0.6 mbar reduced pressure, -52°C). The masses of dried samples were recorded prior to mechanical grinding with a mortar and pestle. Ground algal powder was stirred at 500 rpm in a 95% ethanol solution (1mL solvent: 1g dry algae) for 15 minutes before saponification. Solid sodium hydroxide (30% w/w NaOH: dry algae) was added to the solution, and saponification was carried out under reflux at 80°C for 2.5h. Following reflux, the hot saponification solution was vacuum filtered to remove remaining solid biomass and the filtrate was allowed to cool overnight to form a solution of solid sodium soaps. Deionized water was added to the filtrate to dissolve soaps, and the mixture was washed with hexane to remove unsaponifiable material (carotenoids, sterols, etc.). Sodium soaps were converted to free fatty acids via protonation with 6M HCl (pH<5, stir 1.5h, rt), and the resulting product was extracted into hexane and dried under rotary evaporation. Free fatty acids (FFAs) were esterified under acidic conditions to produce algal biodiesel (1:4 w/v FFA: MeOH, 0.4 eq. H_2_SO_4_, 80°C, 2h). The final reaction mixture was extracted into hexane and washed with methanol to remove trace remaining FFAs. The solvent was removed under the airstream to yield orange liquid biodiesel.

#### Biodiesel analysis

The final biodiesel products were characterized using TLC and GC-MS. Samples of final biodiesel, intermediate free fatty acids, and a reference of *C. pacifica* total lipids prepared from a continuous hexane extraction were analyzed by TLC using glass-backed silica gel 60 plates with a 70/30/1 hexane/diethyl ether/acetic acid eluent. Plates were visualized with a cupric sulfate stain (10% (w/v) CuSO_4_, 4% (v/v) H_2_SO_4_, 4% (v/v) H_3_PO_4_ in MeOH) and charring at 160°C^66^.

GC-MS analysis of biodiesel was performed on an Agilent 7890A GC system connected to an Agilent 5977C GC/MSD. Samples were separated on an Agilent HP-5MS UI 30m x 0.25mm x 0.25um GCMS column with hydrogen as carrier gas and a gradient of 3°C/min from 70°C to 250°C over 60 mins. Integration values from associated peaks were used to determine the relative abundance of different FAMEs comprising each biodiesel sample.

### Conversion of algal biomass to TPUs

The 75% algae-based TPU material was prepared as previously described utilizing a polyester poylol^69,70,85^. The synthesis involved creating a polyester polyol composed entirely of algae-derived aromatic and aliphatic diacids from *C. pacifica* combined with a linear diol. This algae-based polyester polyol was reacted with an algal linear diisocyanate from *C. pacifica*, resulting in the formation of algae TPU, designated as A2141. A2141 had mechanical properties tested in accordance with the American Society for Testing and Materials (ASTM) under tests D2240 and D624-00 for hardness and tensile strength, respectively. Gel permeation chromatography (GPC) was carried out on a Malvern GPC system with a Tosoh TSKgel SuperHZM-N and guard columns with molecular weights and distribution relative to a polystyrene standard. THF served as the polymer solvent and eluent. A2141 was dissolved in DMF at a concentration of 18% and applied to cotton canvas fabric using a coating spreader set to a thickness of 0.05 mm. Water contact angle was tested on a Goniometer, ramé-hart™Model 200, and stiffness was in accordance with ASTM D1388-18.

## Conflict of Interest

**SM** is a founder of and holds equity in Algenesis Inc., a company that could potentially benefit from this research. **MT** is an employee and shareholder in Algenesis Inc. The other authors declare that their research was conducted without any commercial or financial relationships that could be perceived as potential conflicts of interest.

## Authors Contributions

**AG** & **JVDM**: conceptualized the research framework, performed experiments, and contributed to writing, editing, coordinating, and conceptualization of the original draft; **KWF, AB**: performed experiments and contributed to writing, editing, coordinating, and conceptualization of the original draft; **MT**, **KK**, **CD**, **BS, AM**: performed experiments; **SM**: contributed to the writing and editing of the original manuscript and secured financial support for the research.

## Funding

This work was supported by the U.S. Department of Energy’s Office of Energy Efficiency and Renewable Energy (EERE) under the APEX award number DE-EE0009671.

## Acknowledgments

The authors thank Jennifer Santini and Marcy Erb (UCSD School of Medicine Microscopy Core, supported by NINDS P30NS047101 grant) for help with confocal microscopy. The authors extend their sincere gratitude to members of Mayfield Laboratory for their invaluable support. They also wish to thank the United States Department of Energy, the California Center for Algae Biotechnology, and the University of California San Diego for their generous support and contributions.

## References

1. Bongaarts, J. IPCC, 2023: Climate Change 2023: Synthesis Report. IPCC, 184 p., doi: 10.59327/IPCC/AR6-9789291691647. *Popul. Dev. Rev.* **n/a**, (2024).

2. Barrett, C. B. & Lentz, E. C. Food Insecurity. in Oxford Research Encyclopedia of International Studies (2010). doi:10.1093/acrefore/9780190846626.013.438.

3. Ahmed, N., Khan, T. I. & Augustine, A. CLIMATE CHANGE AND ENVIRONMENTAL DEGRADATION: A SERIOUS THREAT TO GLOBAL SECURITY. Eur. J. Soc. Sci. Stud. (2018) doi:10.46827/ejsss.v0i0.394.

4. Foley, J. A. et al. Solutions for a cultivated planet. Nature 478, 337–342 (2011).

5. Owusu, P. A. & Asumadu-Sarkodie, S. A review of renewable energy sources, sustainability issues and climate change mitigation. Cogent Eng. 3, 1167990 (2016).

6. Narodoslawsky, M., Niederl-Schmidinger, A. & Halasz, L. Utilising renewable resources economically: new challenges and chances for process development. J. Clean. Prod. 16, 164– 170 (2008).

7. Østergaard, P. A., Duic, N., Noorollahi, Y., Mikulcic, H. & Kalogirou, S. Sustainable development using renewable energy technology. Renew. Energy 146, 2430–2437 (2020).

8. Zhang, Y.-H. P. Next generation biorefineries will solve the food, biofuels, and environmental trilemma in the energy–food–water nexus. Energy Sci. Eng. 1, 27–41 (2013).

9. Döös, B. R. Population growth and loss of arable land. Glob. Environ. Change 12, 303–311 (2002).

10. Wyman, R. J. The Effects of Population on the Depletion of Fresh Water. Popul. Dev. Rev. 39, 687–704 (2013).

11. Dolganyuk, V. et al. Microalgae: A Promising Source of Valuable Bioproducts. Biomolecules 10, 1153 (2020).

12. Torres-Tiji, Y., Fields, F. J. & Mayfield, S. P. Microalgae as a future food source. Biotechnol. Adv. 41, 107536 (2020).

13. Diaz, C. J. et al. Developing algae as a sustainable food source. Front. Nutr. 9, (2023).

14. Barbosa, M. J., Janssen, M., Südfeld, C., D’Adamo, S. & Wijffels, R. H. Hypes, hopes, and the way forward for microalgal biotechnology. Trends Biotechnol. 41, 452–471 (2023).

15. Khan, M. I., Shin, J. H. & Kim, J. D. The promising future of microalgae: current status, challenges, and optimization of a sustainable and renewable industry for biofuels, feed, and other products. Microb. Cell Factories 17, 36 (2018).

16. Nascimento, I. A. et al. Microalgae Versus Land Crops as Feedstock for Biodiesel: Productivity, Quality, and Standard Compliance. BioEnergy Res. 7, 1002–1013 (2014).

17. Correa, D. F. et al. Freeing land from biofuel production through microalgal cultivation in the Neotropical region. Environ. Res. Lett. 15, 094094 (2020).

18. Ghasemi, Y. et al. Microalgae biofuel potentials (Review). Appl. Biochem. Microbiol. 48, 126– 144 (2012).

19. Hemaiswarya, S., Raja, R., Ravi Kumar, R., Ganesan, V. & Anbazhagan, C. Microalgae: a sustainable feed source for aquaculture. World J. Microbiol. Biotechnol. 27, 1737–1746 (2011).

20. Onen Cinar, S., et al. Bioplastic Production from Microalgae: A Review. Int. J. Environ. Res. Public. Health 17, 3842 (2020).

21. Yan, N., Fan, C., Chen, Y. & Hu, Z. The Potential for Microalgae as Bioreactors to Produce Pharmaceuticals. Int. J. Mol. Sci. 17, 962 (2016).

22. Mourelle, M. L., Gómez, C. P. & Legido, J. L. The Potential Use of Marine Microalgae and Cyanobacteria in Cosmetics and Thalassotherapy. Cosmetics 4, 46 (2017).

23. Rajesh Banu, J., Preethi, Kavitha, S., Gunasekaran, M. & Kumar, G. Microalgae based biorefinery promoting circular bioeconomy-techno economic and life-cycle analysis. Bioresour. Technol. 302, 122822 (2020).

24. Grobbelaar, J. U. Microalgae mass culture: the constraints of scaling-up. J. Appl. Phycol. 24, 315–318 (2012).

25. Dahlin, L. R. et al. Down-Selection and Outdoor Evaluation of Novel, Halotolerant Algal Strains for Winter Cultivation. Front. Plant Sci. 9, (2018).

26. Zhu, Z., Jiang, J. & Fa, Y. Overcoming the Biological Contamination in Microalgae and Cyanobacteria Mass Cultivations for Photosynthetic Biofuel Production. Molecules 25, 5220 (2020).

27. Klein, B., Davis, R. & Wiatrowski, M. Algal Biomass Production via Open Pond Algae Farm Cultivation: 2023 State of Technology and Future Research. https://www.osti.gov/biblio/2370102 (2024) doi:10.2172/2370102.

28. Lafarga, T., Sánchez-Zurano, A., Morillas-España, A. & Acién-Fernández, F. G. Extremophile microalgae as feedstock for high-value carotenoids: A review. Int. J. Food Sci. Technol. 56, 4934–4941 (2021).

29. Sydney, E. B. et al. Biomolecules from extremophile microalgae: From genetics to bioprocessing of a new candidate for large-scale production. Process Biochem. 87, 37–44 (2019).

30. Malavasi, V., Soru, S. & Cao, G. Extremophile Microalgae: the potential for biotechnological application. J. Phycol. 56, 559–573 (2020).

31. Varshney, P., Mikulic, P., Vonshak, A., Beardall, J. & Wangikar, P. P. Extremophilic micro-algae and their potential contribution in biotechnology. Bioresour. Technol. 184, 363–372 (2015).

32. Sproles, A. E. et al. Recent advancements in the genetic engineering of microalgae. Algal Res. 53, 102158 (2021).

33. Gupta, A. et al. Harnessing genetic engineering to drive economic bioproduct production in algae. Front. Bioeng. Biotechnol. 12, (2024).

34. Fajardo, C. et al. Advances and challenges in genetic engineering of microalgae. Rev. Aquac. 12, 365–381 (2020).

35. Maltsev, Y., Maltseva, K., Kulikovskiy, M. & Maltseva, S. Influence of Light Conditions on Microalgae Growth and Content of Lipids, Carotenoids, and Fatty Acid Composition. Biology 10, 1060 (2021).

36. Wong, Y.-K. et al. Effects of Light Intensity, Illumination Cycles on Microalgae Haematococcus Pluvialis for Production of Astaxanthin. J. Mar. Biol. Aquac. 2, (2016).

37. Meagher, E., Rangsrikitphoti, P., Faridi, B., Zamzam, G. & Durnford, D. G. Photoacclimation to high-light stress in *Chlamydomonas reinhardtii* during conditional senescence relies on generating pH-dependent, high-quenching centres. Plant Physiol. Biochem. 158, 136–145 (2021).

38. Wahidin, S., Idris, A. & Shaleh, S. R. M. The influence of light intensity and photoperiod on the growth and lipid content of microalgae *Nannochloropsis* sp. Bioresour. Technol. 129, 7–11 (2013).

39. Hlavova, M., Turoczy, Z. & Bisova, K. Improving microalgae for biotechnology — From genetics to synthetic biology. Biotechnol. Adv. 33, 1194–1203 (2015).

40. Trovão, M. et al. Random Mutagenesis as a Promising Tool for Microalgal Strain Improvement towards Industrial Production. Mar. Drugs 20, 440 (2022).

41. Arora, N., Yen, H.-W. & Philippidis, G. P. Harnessing the Power of Mutagenesis and Adaptive Laboratory Evolution for High Lipid Production by Oleaginous Microalgae and Yeasts. Sustainability 12, 5125 (2020).

42. Fields, F. J., Ostrand, J. T., Tran, M. & Mayfield, S. P. Nuclear genome shuffling significantly increases production of chloroplast-based recombinant protein in Chlamydomonas reinhardtii. Algal Res. 41, 101523 (2019).

43. Ibáñez-Salazar, A. et al. Over-expression of Dof-type transcription factor increases lipid production in Chlamydomonas reinhardtii. J. Biotechnol. 184, 27–38 (2014).

44. Salas-Montantes, C. J. et al. Lipid accumulation during nitrogen and sulfur starvation in Chlamydomonas reinhardtii overexpressing a transcription factor. J. Appl. Phycol. 30, 1721– 1733 (2018).

45. Zhang, J. et al. Overexpression of the soybean transcription factor GmDof4 significantly enhances the lipid content of Chlorella ellipsoidea. Biotechnol. Biofuels 7, 128 (2014).

46. Jia, B. et al. Understanding the functions of endogenous DOF transcript factor in Chlamydomonas reinhardtii. Biotechnol. Biofuels 12, 67 (2019).

47. Jia, B. et al. Increased Lipids in Chlamydomonas reinhardtii by Multiple Regulations of DOF, LACS2, and CIS1. Int. J. Mol. Sci. 23, 10176 (2022).

48. Tokunaga, S., Sanda, S., Uraguchi, Y., Nakagawa, S. & Sawayama, S. Overexpression of the DOF-Type Transcription Factor Enhances Lipid Synthesis in Chlorella vulgaris. Appl. Biochem. Biotechnol. 189, 116–128 (2019).

49. Zheng, Y. et al. iTAK: A Program for Genome-wide Prediction and Classification of Plant Transcription Factors, Transcriptional Regulators, and Protein Kinases. Mol. Plant 9, 1667–1670 (2016).

50. Petroll, R. et al. Signatures of Transcription Factor Evolution and the Secondary Gain of Red Algae Complexity. Genes 12, 1055 (2021).

51. Jin, J. et al. PlantTFDB 4.0: toward a central hub for transcription factors and regulatory interactions in plants. Nucleic Acids Res. 45, D1040–D1045 (2017).

52. Malinova, I. et al. Reduction of the Cytosolic Phosphoglucomutase in Arabidopsis Reveals Impact on Plant Growth, Seed and Root Development, and Carbohydrate Partitioning. PLoS ONE 9, e112468 (2014).

53. Periappuram, C. et al. The Plastidic Phosphoglucomutase from Arabidopsis. A Reversible Enzyme Reaction with an Important Role in Metabolic Control1. Plant Physiol. 122, 1193–1200 (2000).

54. Caspar, T., Huber, S. C. & Somerville, C. Alterations in Growth, Photosynthesis, and Respiration in a Starchless Mutant of Arabidopsis thaliana (L.) Deficient in Chloroplast Phosphoglucomutase Activity 1. Plant Physiol. 79, 11–17 (1985).

55. Hanson, K. R. & McHale, N. A. A Starchless Mutant of Nicotiana sylvestris Containing a Modified Plastid Phosphoglucomutase. Plant Physiol. 88, 838–844 (1988).

56. Harrison, Hedley, & Wang. Evidence that the rug3 locus of pea (Pisum sativum L.) encodes plastidial phosphoglucomutase confirms that the imported substrate for starch synthesis in pea amyloplasts is glucose-6-phosphate. Plant J. 13, (1998).

57. Koo, K. M. et al. The Mechanism of Starch Over-Accumulation in Chlamydomonas reinhardtii High-Starch Mutants Identified by Comparative Transcriptome Analysis. Front. Microbiol. 8, (2017).

58. Takeshita, T., Takeda, K., Ota, S., Yamazaki, T. & Kawano, S. A Simple Method for Measuring the Starch and Lipid Contents in the Cell of Microalgae. Cytologia (Tokyo) 80, 475–481 (2015).

59. Harmon, V. L. et al. Reliability metrics and their management implications for open pond algae cultivation. Algal Res. 55, 102249 (2021).

60. Novoveská, L. et al. Overview and Challenges of Large-Scale Cultivation of Photosynthetic Microalgae and Cyanobacteria. Mar. Drugs 21, 445 (2023).

61. Rawat, I., Ranjith Kumar, R., Mutanda, T. & Bux, F. Biodiesel from microalgae: A critical evaluation from laboratory to large scale production. Appl. Energy 103, 444–467 (2013).

62. Fields, F. J. et al. Annual productivity and lipid composition of native microalgae (Chlorophyta) at a pilot production facility in Southern California. Algal Res. 56, 102307 (2021).

63. Halim, R., Danquah, M. K. & Webley, P. A. Extraction of oil from microalgae for biodiesel production: A review. Biotechnol. Adv. 30, 709–732 (2012).

64. Giorno, F., Mazzei, R. & Giorno, L. Purification of triacylglycerols for biodiesel production from *Nannochloropsis* microalgae by membrane technology. Bioresour. Technol. 140, 172–178 (2013).

65. Fang, Y.-R., Yeh, Y. & Liu, H.-S. A novel strategy of biodiesel production from wet microalgae by direct saponification–esterification conversion (DSEC). J. Taiwan Inst. Chem. Eng. 83, 23–31 (2018).

66. Schoepp, N. G., Wong, W., Mayfield, S. P. & Burkart, M. D. Bulk solvent extraction of biomass slurries using a lipid trap. RSC Adv. 5, 57038–57044 (2015).

67. de Souza, F. M., Kahol, P. K. & Gupta, R. K. Introduction to Polyurethane Chemistry. in Polyurethane Chemistry: Renewable Polyols and Isocyanates vol. 1380 1–24 (American Chemical Society, 2021).

68. Liang, C. et al. Material Flows of Polyurethane in the United States. Environ. Sci. Technol. 55, 14215–14224 (2021).

69. Bruckbauer, A. et al. Renewable and Biodegradable Polyurethane Foams with Aliphatic Diisocyanates. Macromolecules 57, 2879–2887 (2024).

70. Rajput, B. S., Hai, T. A. P. & Burkart, M. D. High Bio-Content Thermoplastic Polyurethanes from Azelaic Acid. Molecules 27, 4885 (2022).

71. Law, K.-Y. Definitions for Hydrophilicity, Hydrophobicity, and Superhydrophobicity: Getting the Basics Right. J. Phys. Chem. Lett. 5, 686–688 (2014).

72. Choi, H.-J., Kim, M. S., Ahn, D., Yeo, S. Y. & Lee, S. Electrical percolation threshold of carbon black in a polymer matrix and its application to antistatic fibre. Sci. Rep. 9, 6338 (2019).

73. Zhu, Z. et al. A Carbon Fixation Enhanced Chlamydomonas reinhardtii Strain for Achieving the Double-Win Between Growth and Biofuel Production Under Non-stressed Conditions. Front. Bioeng. Biotechnol. 8, (2021).

74. Lucker, B. F. et al. Selection-enriched genomic loci (SEGL) reveals genetic loci for environmental adaptation and photosynthetic productivity in *Chlamydomonas reinhardtii*. Algal Res. 64, 102709 (2022).

75. Wilcox, J., Psarras, P. C. & Liguori, S. Assessment of reasonable opportunities for direct air capture. Environ. Res. Lett. 12, 065001 (2017).

76. Barlow, J., Sims, R. C. & Quinn, J. C. Techno-economic and life-cycle assessment of an attached growth algal biorefinery. Bioresour. Technol. 220, 360–368 (2016).

77. Rony, Z. I. et al. Unanswered issues on decarbonizing the aviation industry through the development of sustainable aviation fuel from microalgae. Fuel 334, 126553 (2023).

78. Said, Z. et al. Multi-attribute optimization of sustainable aviation fuel production-process from microalgae source. Fuel 324, 124759 (2022).

79. Izanlou, Z., Akhavan Mahdavi, M., Gheshlaghi, R. & Karimian, A. Sequential extraction of value-added bioproducts from three Chlorella strains using a drying-based combined disruption technique. Bioresour. Bioprocess. 10, 44 (2023).

80. Hammann, W., Ross, A. & Seames, W. Sequential Extraction of Carbohydrates and Lipids from Chlorella vulgaris Using Combined Physical and Chemical Pre-Treatments. ChemEngineering 8, 11 (2024).

81. Findinier, J. Autolysin Production from Chlamydomonas reinhardtii. Bio-Protoc. 13, e4705 (2023).

82. Molino, J. V. D., Carvalho, J. C. M. de & Mayfield, S. P. Comparison of secretory signal peptides for heterologous protein expression in microalgae: Expanding the secretion portfolio for Chlamydomonas reinhardtii. PLOS ONE 13, e0192433 (2018).

83. Bligh, E. G. & Dyer, W. J. A RAPID METHOD OF TOTAL LIPID EXTRACTION AND PURIFICATION. https://cdnsciencepub.com/doi/abs/10.1139/o59-099 (1959).

84. Viegas, C. V. et al. Algal products beyond lipids: Comprehensive characterization of different products in direct saponification of green alga *Chlorella* sp. Algal Res. 11, 156–164 (2015).

85. Rajput, B. S. et al. Renewable low viscosity polyester-polyols for biodegradable thermoplastic polyurethanes. J. Appl. Polym. Sci. 139, e53062 (2022).

