## Supplementary Figures for "Engineering the novel extremophile alga *Chlamydomonas pacifica* for high lipid and high starch production as a path to developing commercially relevant strains"

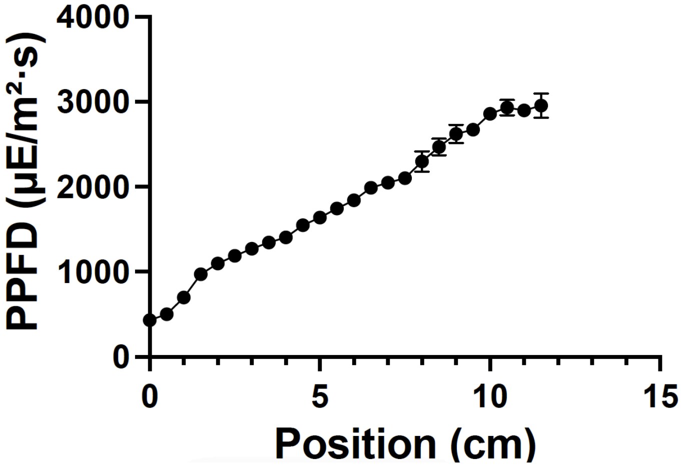


**Supplementary Figure 1**: Light gradient across a culture plate (n=3).

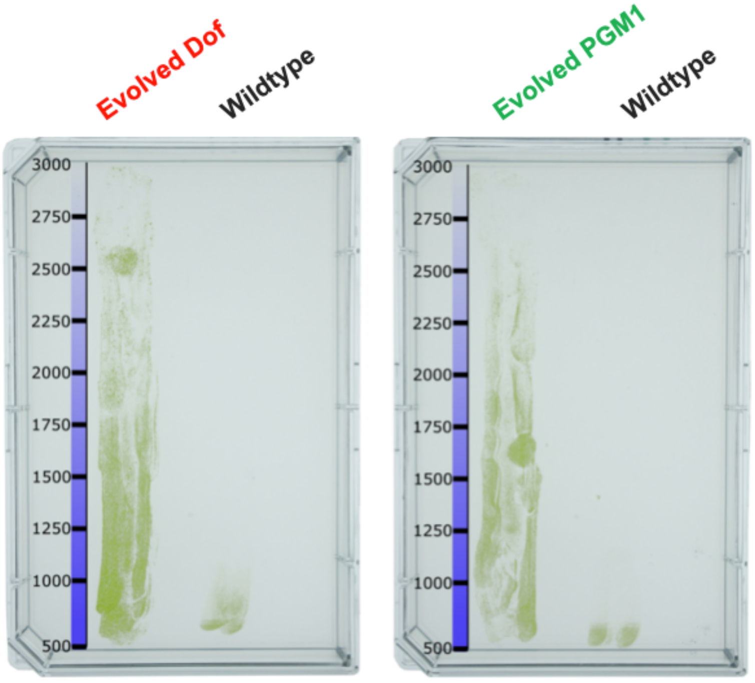


**Supplementary Figure 2: Light intensity tolerance of evolved transgenic strains**. The plates demonstrate the growth of wildtype *C. pacifica* compared to the genetically modified evolved *Dof* and evolved *PGM1*. The unit of light intensity is µE/m²/s.

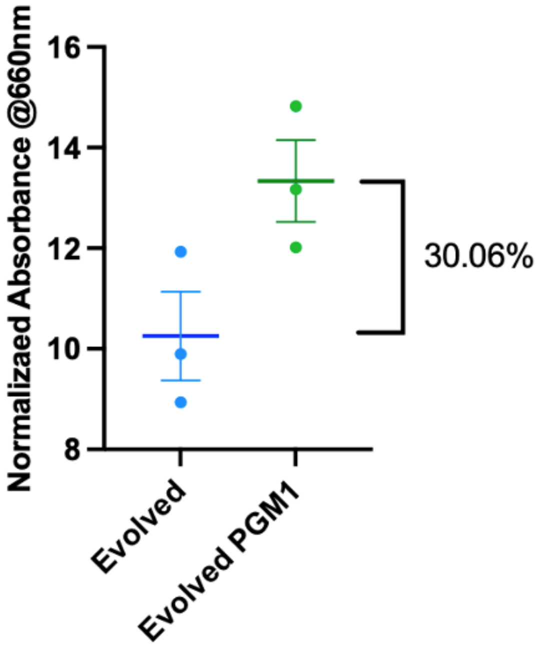


**Supplementary Figure 3: High pH tolerance of wild type and evolved strain**.
Lugol stain absorbance measurements using a plate reader to quantify starch content. Each dot represents a biological replicate.
